# Arc Capsids Facilitate the Transfer of Muscleblind

**DOI:** 10.64898/2026.03.06.708592

**Authors:** Max Zinter, Cong Xiao, Peter M’Angale, Rubing Zhao-Shea, Timothy Freels, Andrew R. Tapper, Travis Thomson

## Abstract

The *Drosophila* activity-regulated cytoskeletal-associated protein (dArc1) can facilitate viral-like synaptic transfer of its own mRNA through dArc1 capsid formation. This transfer promotes synaptic maturation at the *Drosophila* neuromuscular junction and shows conservation to the mammalian neural synapse through the dArc1 mammalian ortholog, Arc. Recently, we established that dArc1 can interact with several transcripts other than its own in *Drosophila* including the transcript of muscleblind (Mbl), an RNA splicing factor known to be involved in neuronal and muscle development. Here, we demonstrate this interaction is further conserved to Arc and the mammalian Mbl ortholog Muscleblind Like Splicing Regulator 1 (Mbnl1). In the mouse neuro2a (N2A) cell line, immunoprecipitation of Arc protein enriches for both the *Arc* and *Mbnl1* transcript. Upon differentiation of N2A cells, the ability of Arc to bind its own transcript and *Mbnl1* are abolished while potassium stimulation of these cells restored Arc interactions with both transcripts, indicating that this interaction is enhanced by neuronal activity. This interaction is further conserved to the mammalian central nervous system, where *Mbnl1* shows increased colocalization with Arc protein in the dentate gyrus of foot-shocked mice. Furthermore, we demonstrate that both *Arc* and *Mbnl1* RNA can be detected in extracellular vesicles (EVs), and that *Mbnl1*, unlike the *Arc* transcript, is not directly encapsulated by Arc protein. We additionally observe MblA crosses the *Drosophila* NMJ, likely within EVs, and postsynaptic MblA accumulation is dependent on presynaptic pools of dArc1. Taken together, our data suggest that Arc protein interacts with *Mbnl1* RNA in an activity-dependent manner and this interaction may facilitate transsynaptic transfer of *Mbnl1* RNA through EVs with implications for neurodevelopment.

**Significance:** The immediate early gene Arc is known to play roles in LTP, LTD, memory and is dysregulated in diseases of the CNS in addition to forming a viral-like capsid to allow for intercellular transfer of its own mRNA in a pathway referred to as the Viral-Like Synaptic Transfer of RNA (ViSyToR). Investigating additional members of the ViSyToR pathway will help increase our understanding of how Arc can play such diverse roles in the CNS. In this study, we establish that transcript for Muscleblind is an additional ViSyToR member in both *Drosophila* and the mammalian CNS.

## Introduction

The immediate early gene Activity-Regulated Cytoskeletal-Associated protein (Arc/Arg3.1) is a known master regulator of synaptic plasticity involved in long-term potentiation (LTP), long-term depression (LTD), learning and memory via its ability to mediate AMPA receptor endocytosis, regulate actin cytoskeleton dynamics and suppress GluA1 transcription[1-4]. This central role in cell-autonomous plasticity has long been the focus of Arc biology. However, the discovery that Arc is a domesticated gene derived from a retrotransposon has unveiled a remarkable, non-cell autonomous function in intercellular communication[5-8].

Both mammalian Arc and its *Drosophila* ortholog, dArc1, possess a domain homologous to the retroviral Group-specific antigen (Gag) protein. This domain enables Arc protein to self-assemble into virus-like capsids that encapsulate their own mRNA. These capsids are then packaged into extracellular vesicles (EVs) and released in a process termed the Viral-Like Synaptic Transfer of RNA (ViSyToR) pathway. This intercellular transfer has been demonstrated to be vital for non-cell autonomous LTD, *Drosophila* sucrose reward valuation, and post-synaptic maturation of the *Drosophila* neuromuscular junction (NMJ[7, 9-11]. Until now, this repurposed viral machinery has been understood primarily as a highly specialized system for Arc to propagate its own genetic material across synapses, leaving open the critical question of whether this pathway could be leveraged by the host cell for the transfer of other, functionally distinct RNAs.

The synapse is a dynamic arena where domesticated and active retroelements interact, creating selective pressures that shape neuronal function. A prime example is the Ty1 retrotransposon *Copia*, which, like dArc1, forms capsids and is transferred across the *Drosophila* NMJ. However, the relationship between *Copia* and dArc1 is antagonistic; *Copia* competes with dArc1 and functions as a negative regulator of synaptic plasticity, with the balance between these two elements dictating the synapse’s structural integrity [12]. The existence of such an antagonistic retroelement suggests an ongoing molecular conflict at the synapse. This context transforms the search for other Arc-interacting RNAs from a simple cataloguing exercise into a targeted search for endogenous host RNAs that may function cooperatively within the ViSyToR pathway.

Our investigation is predicated on a series of key findings that connect the *Drosophila* Arc homolog (dArc1) to the mammalian protein MBNL1. Initially, the discovery that dArc1 can trans-synaptically transfer transcripts other than its own was established through RNA immunoprecipitation experiments that identified the RNA splicing factor Muscleblind (Mbl) as a primary dArc1 interactor [13]. The significance of this finding is underscored by the remarkable evolutionary conservation of the Mbl/MBNL protein family; human MBNL1 not only shares homology but can functionally replace *Drosophila* Mbl, rescuing embryonic lethality in mutant larvae [13-15]. MBNL1 is a crucial regulator of RNA processing, directing the transition from fetal to adult alternative splicing patterns by recognizing YGCY motifs UTRs [14, 16, 17]. Its functional importance is further demonstrated in the pathology of myotonic dystrophy type 1 (DM1), where sequestration of MBNL1 by toxic RNA repeats leads to catastrophic missplicing and muscle defects. Most compellingly, a direct link to Arc’s primary domain of function exists, as MBNL1 knockout mice display a significant reduction in the volume of the hippocampus, a brain region defined by high Arc expression [19]. Therefore, the observed interaction between dArc1 and Mbl at the *Drosophila* NMJ, combined with the high degree of conservation, provides a strong rationale for investigating a conserved mammalian ViSyToR pathway involving Arc and MBNL1.

Here, we test the hypothesis that the ViSyToR pathway is not restricted to self-propagation but functions as a versatile platform for the intercellular transfer of endogenous host regulatory RNAs. We investigate the conserved interaction between Arc and the master splicing regulator Mbnl1 as a model for this expanded functionality. We demonstrate that this interaction is dynamically regulated by neuronal activity in a mammalian system and is conserved in vivo. We further show that both transcripts are packaged into EVs and that MblA transfer across the *Drosophila* NMJ is dependent on presynaptic dArc1. Critically, we uncover a novel mechanism of cargo handling within this system, whereby Arc facilitates the EV-mediated transfer of Mbnl1 mRNA without encapsulating it. These findings fundamentally expand the scope of the ViSyToR pathway, revealing it as a sophisticated mechanism for activity-dependent intercellular communication with profound implications for neurodevelopment and plasticity.

## Results

### The dArc1 interaction with *mbl* is conserved to mammals

We first asked whether the ability for Arc to interact with transcripts other than its own is conserved to mammals. To investigate this possibility, we utilized the mouse Neuro2a (N2A) neuroblastoma cell line. We began by immunoprecipitating Arc and then analysing both *arc* and *mbnl1* RNA content by digital PCR (Fig 1A). Our immunoprecipitation approach showed specificity for Arc, as immunoprecipitated product showed an enrichment of Arc when compared to the pre-immune serum and an absence of GAPDH (Fig 1B). Consistent with previous results in *Drosophila*, Arc transcript co-immunoprecipitated with Arc protein (unpaired two-tailed *t* test [*t*_4.187_ =4, *P*=0.0138]; Fig 1C). Interestingly, immunoprecipitation of Arc also showed enrichment for the *mbnl1* transcript, demonstrating Arc’s ability to interact with transcripts other than its own likely being conserved across species (unpaired two-tailed *t* test [*t*_8.141_=4, *P*=0.0012]; Fig 1D).

**Figure 1.**
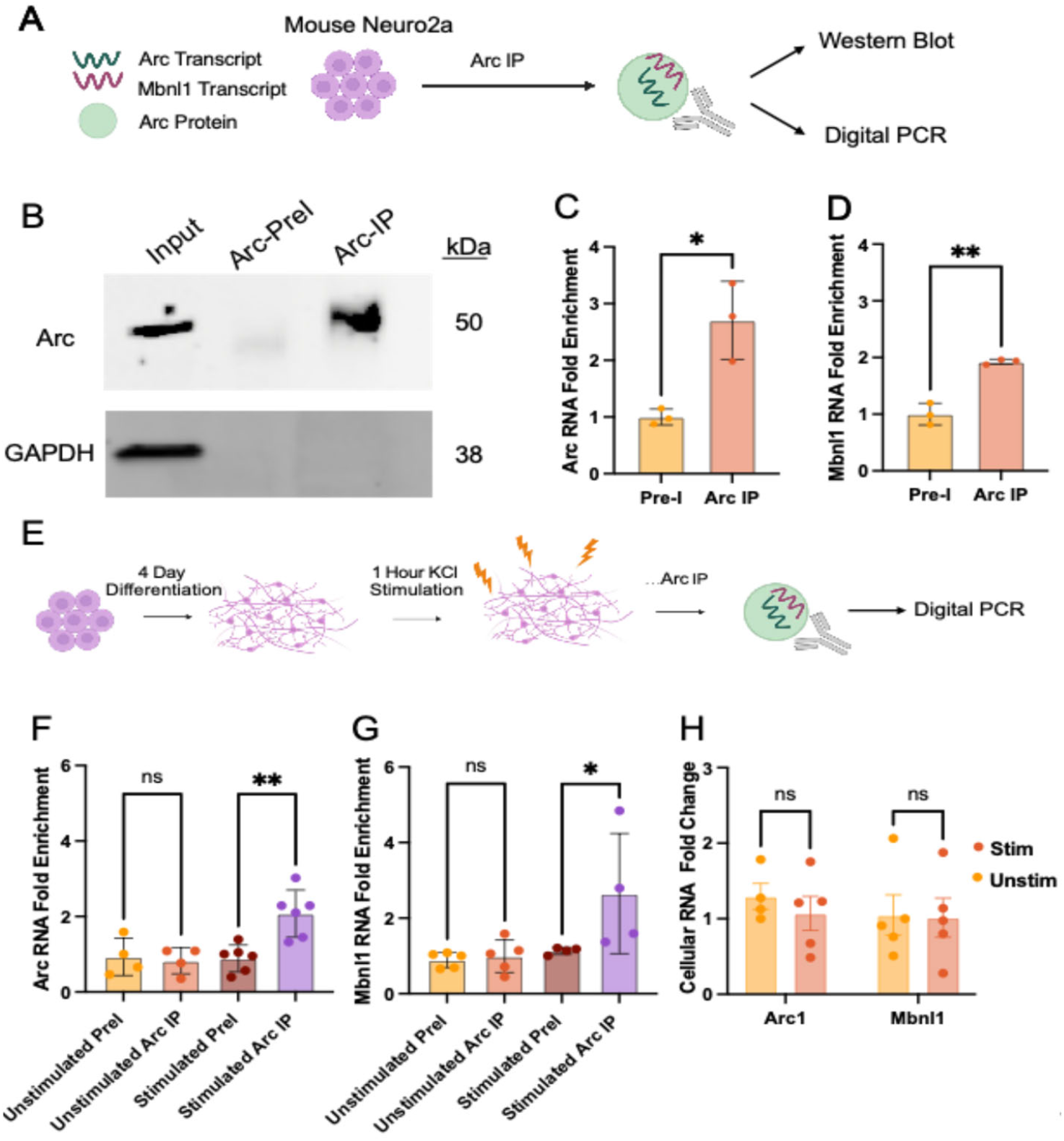
The interaction between dArc1 and *mbl* is conserved to mammals. **A**. Experimental workflow schematic. Arc protein was immunoprecipitated from mouse Neuro2A cells. IP purity was assessed by western blot and dPCR was used to detect Arc and Mbnl1 transcript. **B**. Western blot analysis of protein collected from Neuro2A input, rabbit serum control (Arc-PreI) and immunoprecipitated samples (Arc-IP). **C**. Digital PCR analysis of Arc transcript from pre-immune and immunoprecipitated conditions. **D**. Digital PCR analysis of Mbnl1 transcript from pre-immune and immunoprecipitated conditions. Error bars, mean ± SEM; Statistical analysis conducted using an unpaired T-test; P values: *<0.05, **<0.01. **E**. Experimental workflow schematic. Neuro2A were differentiated in reduced serum media supplemented with 20 μM retinoic acid for 4 days and then stimulated for 1 h with 53 mM KCl stimulation buffer. Arc was immunoprecipitated from cell lysate and Arc and Mbnl1 transcript were detected by dPCR **F**. Digital PCR analysis of Arc transcript from preimmune (Pre-I) and immunoprecipitated (IP) RNA from unstimulated and stimulated differentiated Neuro2A cultures. **G**. Same as in F. for Mbnl1 transcripts. **H**. Digital PCR analysis of Arc and Mbnl1 transcript from cell lysate of unstimulated and stimulated differentiated Neuro2A cultures. Error bars, mean ± SEM; Statistical analysis conducted using a 2-way ANOVA with Bonferroni’s posttests; P values: *<0.05, **<0.01, NS, not significant.

### The Arc-*mbnl1* interaction is dependent on neuronal activity

Given Arc is an immediate early gene and its capsid formation and extracellular release is an activity dependent process [7, 8], we next asked whether Arc’s association with *mbnl1* is similarly dependent on neuronal activity. To this end, we utilized the ability of N2A cells to differentiate into a more neuronal-like state [18]. N2A cells were exposed to reduced serum conditions (2% FBS) and 20 μM retinoic acid for 4 days (Fig 1E, S1A) which produced an increase in neurite-like outgrowths (unpaired two-tailed *t* test [*t*_4.544_=5, *P*=0.0.0061]; Fig S1C), neurite length (unpaired two-tailed *t* test [*t*_8.683_ =8, *P*=0.0001]; Fig S1B) and reduced cell division (unpaired two-tailed *t* test [*t*_7.776_ =8, *P*=0.0001]; Fig S1D) consistent with a more terminally differentiated neuronal-like state. After 4 days of differentiation, cells were then exposed to 53 mM KCl buffer for 1 h to induce membrane depolarization. After 1 h of depolarization, cells and media were collected for immunoprecipitation (Fig 1E). Two-way ANOVA of differentiation revealed a stimulation x *arc* transcript enrichment interaction ([*F*_3,11_=7.797, *P*=0.004]; Fig 1F).

Posttests revealed that, after N2A differentiation Arc no longer co-immunoprecipitated its own transcript (differentiated unstimulated Arc-IP versus differentiated unstimulated pre-immune, *P*=>0.9999; Bonferroni’s posttests; Fig 1F) but stimulation of differentiated N2A restored Arc’s ability to co-immunoprecipitate its own transcript (differentiated stimulated Arc-IP vs differentiated stimulated pre-immune, *P*=0.0031; Bonferroni’s posttests; Fig 1F). Additionally, two-way ANOVA of differentiation revealed a stimulation x *mbnl1* transcript enrichment interaction as well ([*F*_3,10_=4.315, *P*=0.0339]; Fig 1G). Posttests revealed that, differentiation abolished Arc’s ability to co-immunoprecipitate *mbnl1* (differentiated unstimulated Arc-IP versus unstimulated pre-immune, *P*=>0.9999; Bonferroni’s posttests; Fig 1G). Stimulation of differentiated N2A also restored Arc’s ability to co-immunoprecipitate *mbnl1* (differentiated stimulated Arc-IP vs differentiated stimulated pre-immune, *P*=0.0434; Bonferroni’s posttests; Fig 1G). These data suggest the Arc-*mbnl1* interaction is an activity dependent process. This increased interaction was not the result of an increase in *arc* nor *mbnl1* transcription after potassium stimulation, indicating this interaction is likely post transcriptionally regulated (two-way ANOVA; ([*F*_1,15_=0.2855, *P*=0.6010]; Fig 1H).

### The Arc-*mbnl1* interaction is further conserved to the mouse dentate gyrus

Having demonstrated dArc1’s interaction with *mbl* is conserved in a mouse cell line and is likely an activity dependent process, we next asked whether this interaction was further conserved to the mammalian brain. The hippocampus is a known hub for Arc activity in response to learning and memory [19, 20], potentially providing a likely region in the brain to observe this interaction. To test our hypothesis, we administered foot shocks to 6-8 week old mice and collected hippocampal slices for analysis by confocal microscopy. Hippocampal slices where then probed for Arc protein and *mbnl1* transcript by dual immunofluorescence and *in situ* hybridization (Fig 2A). The dentate gyrus showed the greatest density of *mbnl1* (data not shown) and was selected for colocalization analysis. The number of Arc expressing cells in the dentate gyrus from foot-shocked mice was significantly increased compared to dentate gyrus from control mice (Fig 2B2-B2’, unpaired two-tailed *t* test [*t*_4.435_=4, *P*=0.0114]; Fig 2C) while *mbnl1* expression did not differ between groups (Fig 2B1-B1’, unpaired two-tailed [*t* test: *t*_0.8887_=4, *P*=0.4244];; Fig 2D). We next quantified the colocalization of *mbnl1* positive puncta with Arc positive puncta in hippocampus from foot-shocked and control mice. In the dentate gyrus from control mice, we observed a mild positive correlation between *mbnl1* and Arc colocalization. However, dentate gyrus from foot-shocked mice showed a significant increase in correlation coefficient between *mbnl1* and Arc compared to control mice (Fig 2B3-B4’, unpaired two-tailed *t* test [*t*_3.521_ =4, *P*=0.0244]; Fig 3E). These data indicate the activity-dependent interaction between Arc and *mbnl1* is further conserved to the mammalian dentate gyrus.

**Figure 2.**
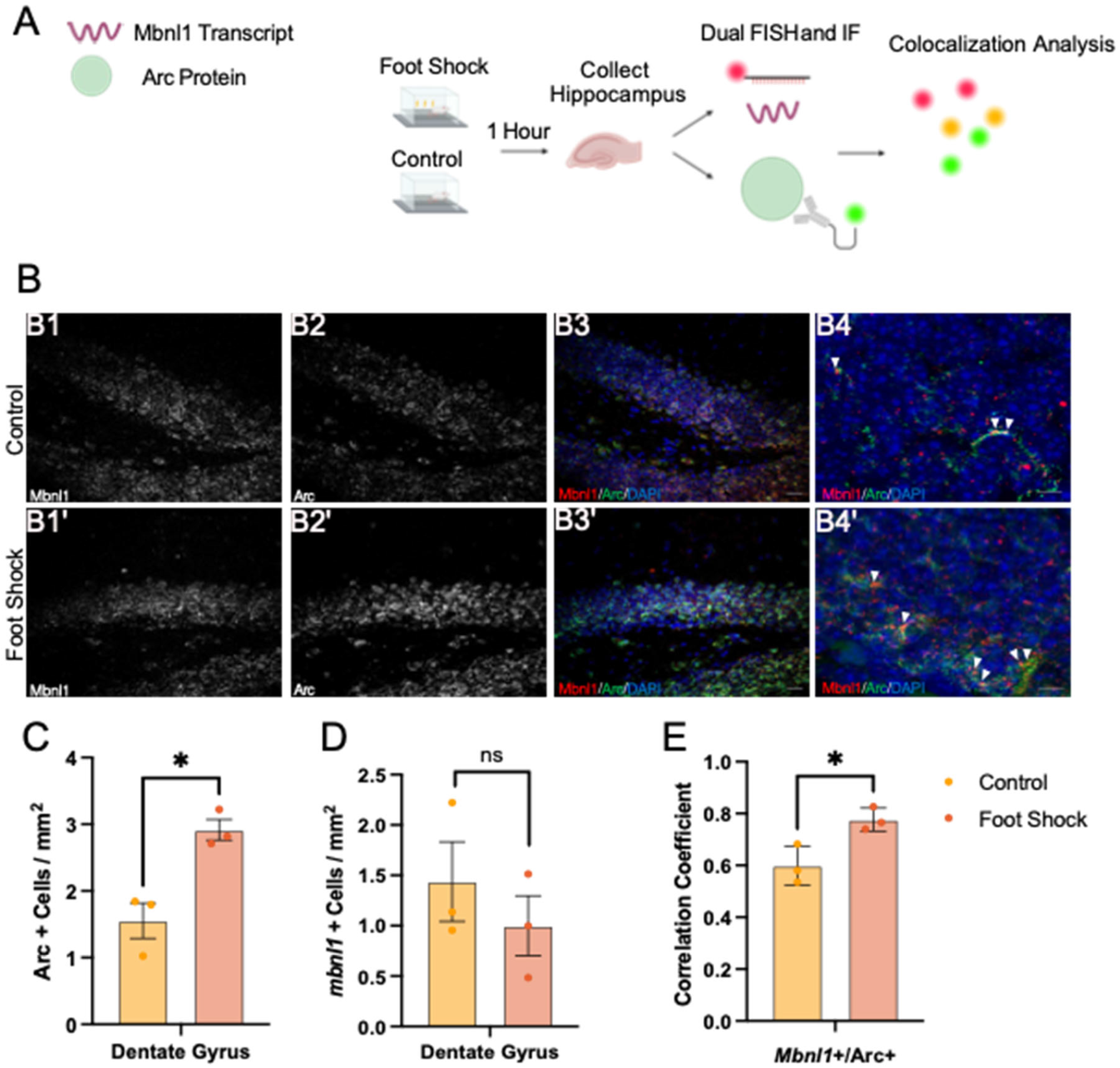
Arc and *mbnl1* colocalize at the mouse dentate gyrus in response to foot shock. **A**. Experimental workflow schematic. Foot shock and control mouse hippocampus was collected 1 hour after foot shock or mock procedure and then dual stained for *mbnl1* and Arc protein prior to colocalization analysis. **B1-B3’**. Representative of micrographs of control and foot shock mouse hippocampus. Scale bar = 20 μm. **B4-B4’** High magnification representative micrographs with arrows marking colocalization events. Scale bar = 5 μm. **C**. Quantification of Arc+ cells **D**. *mbnl1+* cells and **E**. change in *mbnl1* and Arc colocalization quantified by Pearson correlation. Error bars, mean ± SEM; Statistical analysis conducted using an unpaired T-test. P values: *<0.05, NS, not significant.

**Figure 3.**
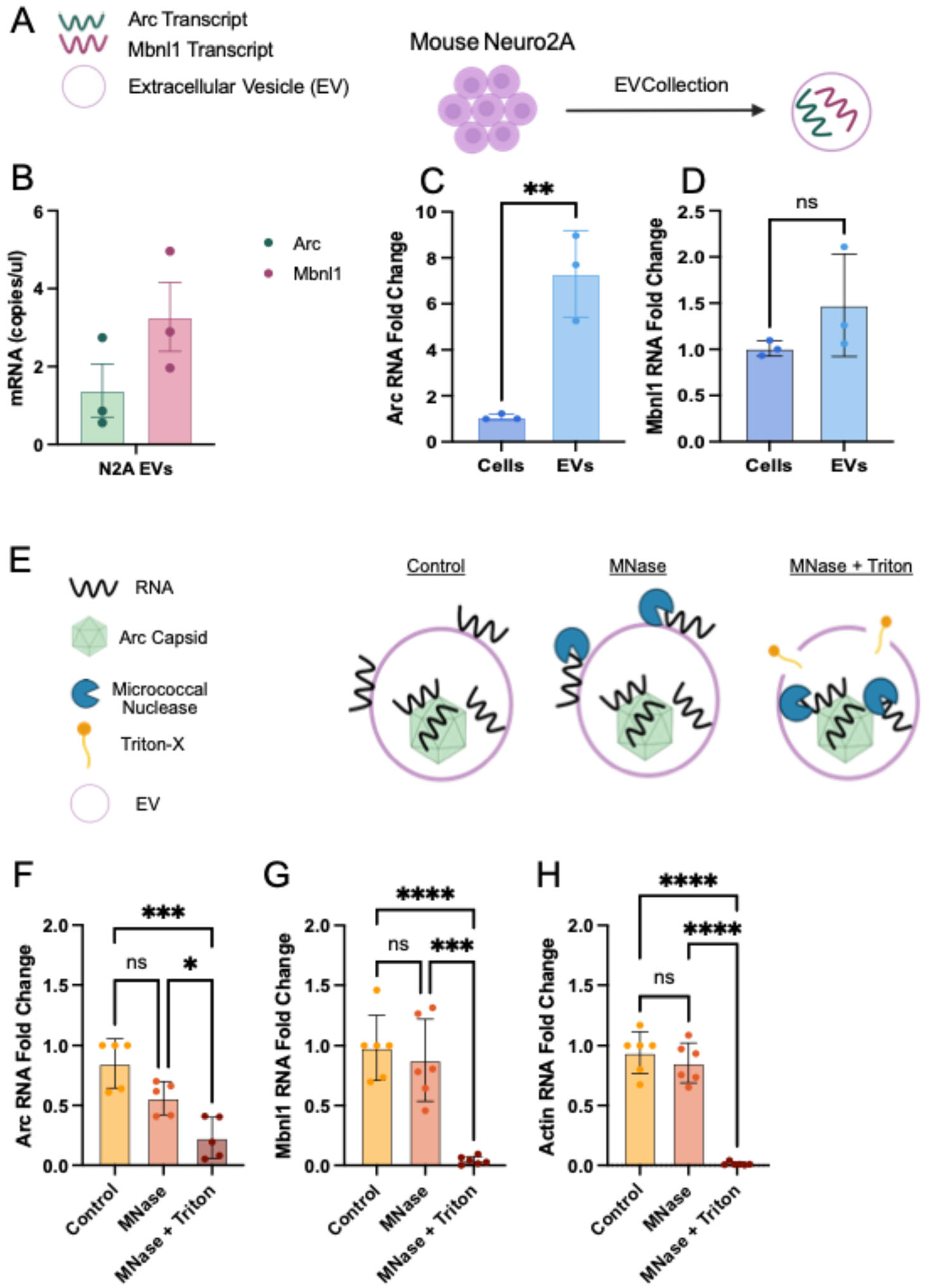
*Mbnl1* is present in mammalian derived extracellular vesicles but not encapsulated by Arc. **A**. Experimental workflow schematic. Extracellular vesicles were isolated from Neuro2A cell supernatant and EV RNA was extracted for downstream analysis. **B**. dPCR detecting Arc and Mbnl1 mRNA present in EVs. **C**. dPCR Analysis of Arc RNA enrichment in EVs compared to cell lysate **D**. Same as an **C** for Mbnl1. Error bars, mean ± SEM; Statistical analysis conducted using an unpaired T test; P values: *<0.05, **<0.01, ***<0.005, NS, not significant. **E**.Experimental workflow schematic. EV derived nucleic acid was sequentially exposed to MNAse digestion to detect protected nucleic acids. Control received no MNAse, MNase received MNase alone and MNase +Triton received MNase paired with detergent to disrupt EV membrane. **F-H** dPCR detection of **F**. Arc, **G**. Mbnl1 and **H**. Actin mRNA after MNase exposure as described in **E**. Error bars, mean ± SEM; Statistical analysis conducted using ANOVA with Bonferroni’s posttest; P values: *<0.05, **<0.01, ***<0.005, ****<0.0001, NS, not significant.

### Arc and mbnl1 transcripts are present in Neuro2A derived extracellular vesicles

Given both dArc1 and Arc have been detected in EVs and EV trafficking is the proposed mechanism of *darc1*/*arc* transfer [7, 8], we next investigated whether *mbnl1* could similarly be detected in EVs. We began by evaluating our ability to isolate a pure population of EVs from N2A supernatant (Fig 3A). Quantification of EV size by tuneable resistive pulse sensing (TRPS) showed a characteristic EV size distribution with a peak at ∼80 nm and transmission electron microscopy demonstrated the canonical cup shape morphology expected from EVs (Fig S1E,S1E inset). Furthermore, EVs show canonical intraluminal markers of EVs (TSG101, HSP70), surface markers (CD81) and are free from organelle contamination (Calnexin) as detected by western blot (Fig S1F). We next evaluated EVs for *arc* and *mbnl1* content by digital PCR. Interestingly, both *arc* and *mbnl1* were detected within N2A derived EVs (Fig 3B). Furthermore, *arc* showed a ∼7 fold enrichment in EVs when compared to cellular *arc* ([unpaired two-tailed *t* test [*t*_5.717_=4, *P*=0.0.0046]; Fig 3C). Interestingly, *mbnl1* did not show any significant enrichment in EVs when compared to cellular derived *mbnl1* ([unpaired two-tailed *t* test [*t*_1.444_=4, *P*=0.2222]; (Fig 3D).

Given the detection of *mbnl1* in N2A derived EVs and knowing Arc’s ability to encapsulate its own transcript, we next asked whether Arc is also capable of encapsulating the *mbnl1* transcript. N2A EVs were collected then either exposed to micrococcal nuclease to digest any nucleic acid riding on the EV surface or micrococcal nuclease paired with Triton-X detergent to disrupt the EV membrane and expose intraluminal RNA cargo. Subsequently, *arc, mbnl1* and *actin* (as a control for EV membrane integrity) were detected by digital PCR and compared against an unexposed EV control condition (Fig 3E). *Arc* was reduced, but still detectable, by exposure to both micrococcal nuclease and triton-X but not micrococcal nuclease alone (one-way ANOVA [*F*_2,12_=15.9, *P*=0.0004]; posttests: nuclease alone versus control, *P* = 0.0670; nuclease and detergent versus control, *P*= 0.0003; nuclease alone versus nuclease and detergent, *P*=0.0325 (Fig 3F)). *Mbnl1* was similarly resistant to nuclease treatment alone but was completely abolished by nuclease and detergent treatment (one-way ANOVA [*F*_2,15_=24.83 *P*=0.0001]; posttests: nuclease alone versus control, *P* = >0.9999; nuclease and detergent versus control, *P*= 0.0001; nuclease alone versus nuclease and detergent, *P*=<0.0001 (Fig 3G)). *Actin*, like *mbnl1*, was resistant to nuclease treatment alone but completely abolished by nuclease and detergent treatment (one-way ANOVA [*F*_2,15_=80.69 *P*=<0.0001]; posttests: nuclease alone versus control, *P* = 0.9271; nuclease and detergent versus control, *P*= <0.0001; nuclease alone versus nuclease and detergent, *P*=<0.0001 (Fig 3H)). These data suggest that all three transcripts are present intraluminally as opposed to on the EV surface. However, *Arc* was partially protected from micrococcal nuclease digestion after Triton-X exposure, suggesting that Arc encapsulates its own transcript but not that of *mbnl1* nor *actin*.

### Potassium stimulation alters neuro2A EV cargo

We next sought to ask whether the EV derived *arc* and *mbnl1* transcripts were influenced by neuronal activity similarly to the interaction between Arc protein and the *mbnl1* transcript. To answer this question, we utilized the 4-day neuro2A differentiation protocol and isolated EVs from the differentiated cell culture media as previously described (Fig S2A). We observe an enrichment of EV derived *arc* when compared to cellular *arc* in both the unstimulated (unpaired two-tailed *t* test [*t*_2.996_=8, *P*=0.0172]; S1B) and stimulated (unpaired two-tailed *t* test [*t*_2.550_=8, *P*=0.0341; S1C). However, *mbnl1* was only enriched in EVs when compared to cell lysate in the unstimulated condition (unpaired two-tailed *t* test [*t*_3.142_=10, *P*=0.0105]; S1D) and was no longer enriched in EVs after stimulation (unpaired two-tailed *t* test [*t*_1.988_=10, *P*=0.0749]; S1E). This result demonstrates neuronal stimulation reduces the amount of EV *mbnl1* but not *arc*.

### Postsynaptic accumulation of *Drosophila* MblA is dependent on presynaptic pools of dArc1

EVs are a proposed vehicle for intercellular communication regulating numerous aspects of synaptic development at the *Drosophila* NMJ and dArc1 has been shown to regulate maturation at the NMJ through the synaptic transfer of EVs [7, 21]. Given this and our data indicating Mbnl1 and Arc transcripts are present in EVs from Neuro2A cells, we used the genetic tractability and compartmentalization of the *Drosophila* NMJ (Fig S2F) to ask whether the presynaptic pool of dArc1 could influence postsynaptic accumulation of MblA, the Mbl isoform shown to be most highly expressed at the NMJ [13]. To this end, we utilized the Gal4-UAS system in *Drosophila* to express a dArc1 RNAi exclusively in the presynaptic neuron using the C380 driver (Fig 4A1-B3). We first validated the dArc1 RNAi line by dPCR, where we observed a significant reduction in both presynaptic (unpaired two-tailed *t* test [*t*_4.635_=4, *P*=0.0098]; Fig 4C) and postsynaptic (unpaired two-tailed *t* test [*t*_7.488_=4, *P*=0.0017]; Fig 4D) *darc1* consistent with previous work [7]. We then assessed accumulation of MblA at the NMJ, we observed no changes in presynaptic MblA accumulation when dArc1 was reduced presynaptically (unpaired two-tailed *t* test [*t*_0.03108_=10, *P*=0.9758]; Fig 4E). Interestingly, MblA was reduced at the postsynaptic muscle (unpaired two-tailed *t* test [*t*_2.597_=10, *P*=0.0266]; Fig 4F) when presynaptic dArc1 was reduced. These data suggest MblA expression at the postsynaptic muscle is dependent on a presynaptic pool of dArc1.

**Figure 4.**
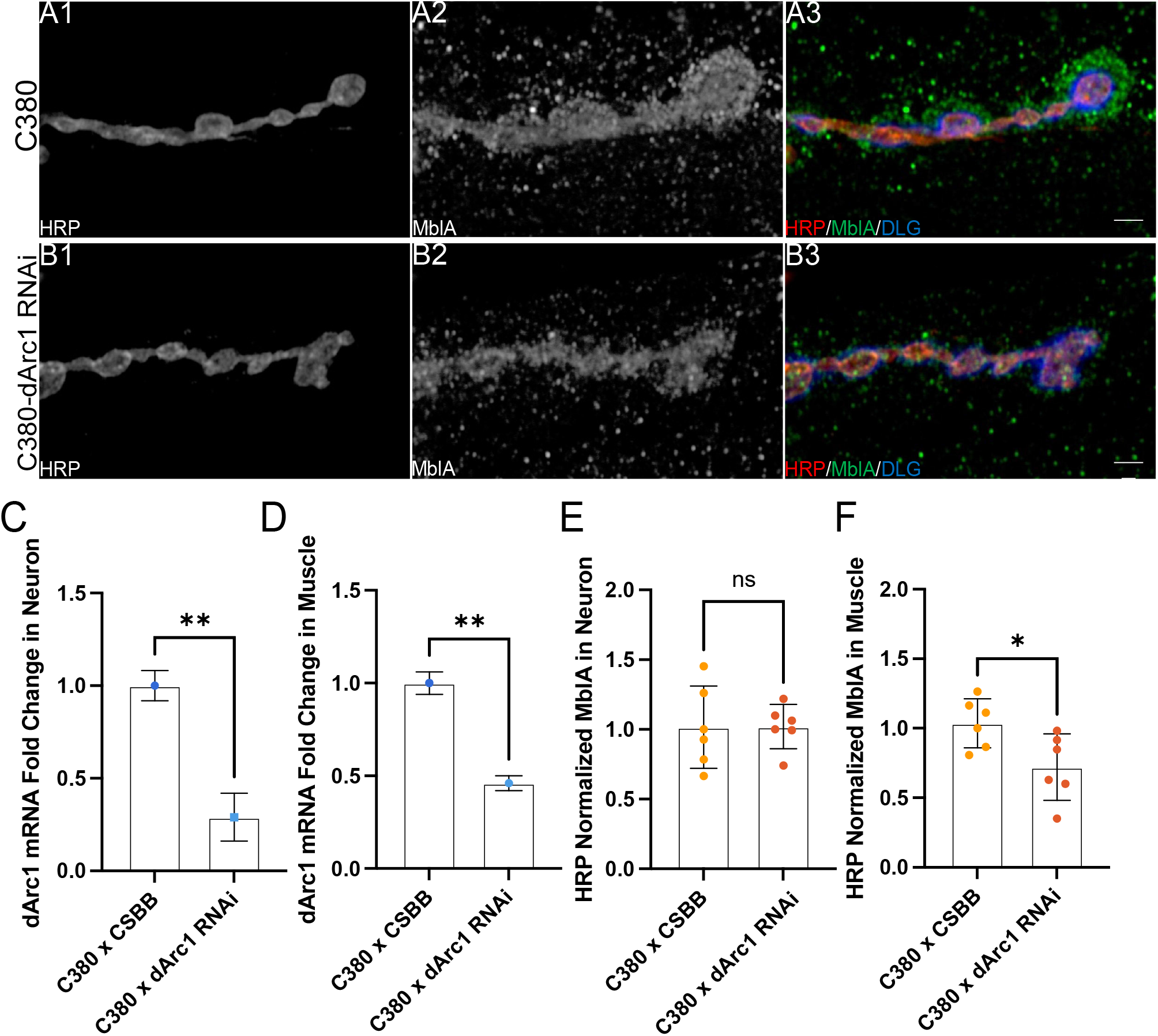
Postsynaptic MblA is dependent on Presynaptic dArc1. **A1-A3**. Representative micrographs of C380 x CSBB 3^rd^ instar larval NMJs stained for **A1**.HRP to mark the presynaptic neuron **A2**. MblA and **A3**. Merged image including DLG to mark the subsynaptic reticulum. **B1-B3**. Representative micrographs of C380 x dArc1 RNAi 3^rd^ instar larval NMJs stained for **B1**.HRP, **B2**. MblA and **B3** merged image including DLG. scale bar = 5 μm **C**. Quantification of dArc1 RNA in the neurons of C380 x CSBB and C380 x dArc1 RNAi flies. **D**. Same as in **C**. but for dArc1 in muscle. **E**. Quantified MblA protein normalized to HRP volume in the presynaptic neuron **F**. Quantified MblA protein normalized to post synaptic volume in the muscle. Error bars, mean ± SEM; Statistical analysis conducted using an unpaired T test; P values: *<0.05, **<0.01, NS, not significant.

### MblA crosses the NMJ synapse and postsynaptic accumulation of MblA is decreased in response to disruption of presynaptic endosomal cycling

We next sought to test whether this postsynaptic pool of MblA could be influenced by the transfer of EVs. We first asked whether MblA itself was capable of transferring across the synapse from the presynaptic neuron to the postsynaptic muscle. To do so, we again utilized the C380 driver to express either a presynaptic GFP or GFP N-terminally tagged to the MblA protein. We then assessed post-synaptic accumulation of GFP signal (Fig 5A1-B4). When GFP alone is driven in the presynaptic neuron, we observe an accumulation of GFP signal within the neuron but not in the postsynaptic muscle indicating GFP alone is incapable of transferring across the synapse (Fig 5A1-A4). When GFP was N-terminally tagged to MblA, we observe a postsynaptic accumulation of MblA in the muscle consistent with MblA being capable of transferring across the synapse (Fig 5B1-B4). To further test whether this could be an EV mediated phenomenon, we expressed a presynaptic Rab11 dominant negative mutant (DN) to reduce EV release at the NMJ. In the Rab11 DN condition, we observe a reduction (unpaired two-tailed *t* test [*t*_3.595_=4, *P*=0.0229]; Fig 5E) in the postsynaptic accumulation of MblA (Fig 5C-E). Taken together, these data indicate MblA is capable of transferring from the presynaptic motoneuron to the postsynaptic muscle and this transfer is likely occurring through EVs.

**Figure 5.**
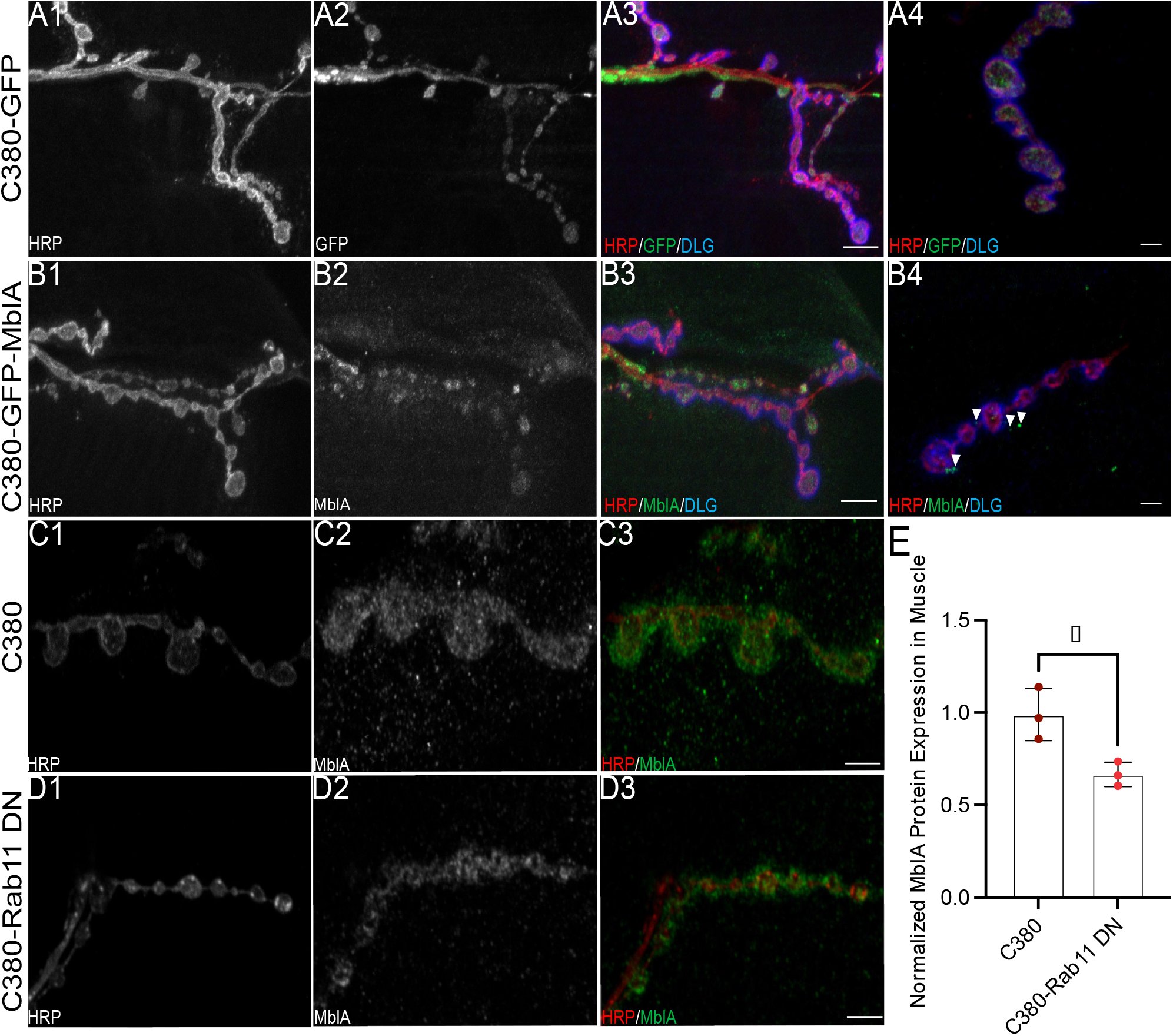
MblA crosses the NMJ likely within EVs. **A**. Micrographs from presynaptically driven GFP 3^rd^ instar larval NMJs stained for **A1**. HRP to mark the presynaptic neuron **A2**. GFP **A3**. merged image including DLG to mark the subsynaptic reticulum, scale bar = 20 μm and **A4**. high magnification merged image, scale bar = 2 μm **B** Micrographs from presynaptically driven GFP-MblA 3^rd^ instar larval NMJs stained for **B1**. HRP **B2**. GFP-MblA **B3**. merged image including DLG, scale bar = 20 μm and **B4**. High magnification merged image, arrows denote postsynaptic accumulation of GFP-MblA, scale bar = 2 μm. **C**. Representative images from C380 x CSBB control 3^rd^ instar larval NMJs stained for **C1**. HRP **C2**. MblA and **C3**. Merged image, scale bar = 5 μm. **D**. Representative images from presynaptically driven Rab11 dominant negative stained for **D1**. HRP **D2**. MblA and **D3**. merged image, scale bar = 5 μm. **E**. Quantification of MblA in the postsynapse normalized to postsynaptic volume. Error bars, mean ± SEM; Statistical analysis conducted using an unpaired T-test; P-values: *<0.05.

## Discussion

Here, we identify the transcript for the splicing factor Mbnl1 as the first endogenous host regulatory mRNA in the ViSyToR pathway and establish the conservation of this expanded cargo capacity in a mammalian system. This discovery is significant as it demonstrates the pathway is not limited to trafficking retrotransposon-derived elements. Furthermore, we uncover an entirely new mechanism for cargo loading within this system. While Arc encapsulates its own mRNA in a protective capsid, our data reveal that Mbnl1 transfer is also Arc-dependent but occurs via a distinct, non-encapsulation mechanism. This suggests a sophisticated dual function for Arc protein: it acts as both a ‘container’ for its own transcript and as a ‘recruiter’ for other regulatory RNAs. This discovery fundamentally expands the scope of the ViSyToR pathway, recasting it as a versatile platform for intercellular communication capable of utilizing multiple molecular strategies to transport distinct classes of RNA cargo.

Our results establish that the physical association between Arc protein and Mbln1 mRNA is a feature conserved from Drosophila to mammals and is dynamically regulated by neuronal activity. In differentiated mouse neuroblastoma cells, which serve as a post-mitotic neuron-like model, this interaction is quiescent but is robustly triggered by membrane depolarization. This finding was validated in vivo, where foot-shock-induced neuronal activation significantly increased the colocalization of Arc protein and Mbnl1 mRNA in the dentate gyrus. This tight coupling to neuronal activity links the ViSyToR pathway directly to the core physiological processes of learning and memory, where Arc is a known master regulator [1-4]. The observation that Arc interacts with Mbnl1 in undifferentiated, proliferating cells suggests a potential role in developmental plasticity that becomes gated by synaptic activity in mature circuits.

Critically, our work elucidates a bimodal loading mechanism for EV cargo sorting, evidenced by our nuclease protection assays. Our data delineate two distinct modes of RNA handling for EV export. The first is the established mechanism of direct encapsulation: consistent with prior work, we show that Arc mRNA within EVs is partially resistant to nuclease digestion even after membrane disruption, confirming that Arc protein assembles into a protective, virus-like capsid around its own transcript. The second, novel mechanism is capsid-independent recruitment. In stark contrast to Arc mRNA, we found that Mbnl1 mRNA, while protected inside EVs, is completely degraded upon membrane lysis. This demonstrates that Mbnl1 is not encapsulated within the Arc capsid. Because the transfer of MblA in *Drosophila* is dependent on presynaptic dArc1, we hypothesize that Arc protein may play a conserved, active role in loading *Mbnl1* mRNA into EVs. While mammalian Arc lacks the NC domain that facilitates RNA binding in dArc1, recent work has demonstrated putative RNA interaction motifs formed after Arc dimerization [22, 23]. This raises the possibility that Arc mediates RNA interactions to recruit cargo into the capsid, a mechanism previously established for dArc1 [7].

How Arc facilitates the loading of non-encapsulated Mbnl1 mRNA into EVs remains an open question. Recent work has shown that Arc capsid assembly and secretion occurs via the endosomal-multivesicular body (MVB) pathway, with Arc protein accumulating within intraluminal vesicles [24]. This localization places Arc at the site of exosome biogenesis, which is known to involve the ESCRT machinery for sorting cargo [7, 24]. It is plausible that Arc acts as a molecular adaptor in this context. The sorting of RNA into EVs is generally an active process mediated by RNA-binding proteins (RBPs) that recognize specific cargo and guide it to the MVB [25, 26]. We propose that Arc may recruit an Mbnl1-RNP complex to the site of ILV formation, thereby facilitating its inclusion into the forming vesicle without direct encapsulation. Although cell type-specific targeting of dArc1 EVs has been previously observed [7] and the molecular mechanisms driving this specificity have recently been elucidated [27], it has not been directly shown that dArc1 and mblA are co-packaged within the same EV. Future work using fluorescently activated vesicle sorting or electron microscopy in situ hybridization approaches should be used to investigate this possibility and further elucidate the mechanisms of this complex pathway.

Our observations in Drosophila provide strong in vivo support for this model of intercellular transfer. We found that postsynaptic accumulation of MblA at the NMJ was dependent on a presynaptic pool of dArc1 and that MblA was capable of transferring across the synapse, likely through EVs. These results are consistent with a role for dArc1 in mediating the loading of MblA into EVs. It remains unclear, however, whether EV-derived mblA is translated after uptake by the postsynaptic muscle. In vitro transcription assays from both Drosophila Schneider 2 and mouse Neuro2A derived EVs would be highly informative as to whether loading of mblA/mbnl1 is an active process intended to provide recipient cells with a supply of MblA/Mbnl1 protein or a method of efficiently degrading transcript that is no longer required.

The functional implications of this expanded ViSyToR pathway are profound. By transferring the mRNA for a master splicing regulator like Mbnl1, a presynaptic neuron could remotely and dynamically alter the alternative splicing landscape of a postsynaptic cell in an activity-dependent manner. This represents a highly sophisticated form of intercellular communication, where one neuron transfers not just a signal, but the molecular machinery to reprogram gene expression in another. Such a mechanism could drive the coordinated, long-lasting changes in the postsynaptic proteome that underpin enduring forms of synaptic plasticity and neurodevelopment.

In conclusion, this work fundamentally expands our understanding of the ViSyToR pathway by demonstrating Arc-dependent packaging and transfer of a non-Arc regulatory RNA, Mbnl1. These data indicate that viral-like transfer of Arc across the synapse is not just for Arc self-propagation; rather the ViSyToR pathway represents a versatile, dual-mechanism system for more sophisticated intercellular communication. This discovery not only provides a new framework for investigating activity-dependent plasticity in the nervous system but also introduces a novel principle of cargo handling in EV biology that is likely to have broad relevance.

## Materials and Methods

### Cell Culture

Mouse Neuro2A cells were cultured in DMEM/F-12 (Life Technologies #11320033) in 10 % exosome-depleted Fetal Bovine Serum (System Biosciences #EXO-FBS) and 1% Penicillin-Streptomycin (Gibco #15140122). Neuro2A were differentiated by seeding 40,000 cells per cm^2^ in 10% FBS and 1% P/S. 24 hours after seeding cells were placed on differentiation media comprised of DMEM/F-12 with 2% exosome-depleted FBS, 20 μM Retinoic Acid (Sigma #R2500) and 1% P/S for four days. For stimulation experiments, differentiated Neuro2A were then exposed to stimulation buffer (53 mM KCl, 3.33 mM HEPES, 0.67 mM CaCl2, 0.33 mM MgCl2, pH=7.4) or PBS control in differentiation media for 1 hour. Both cells and media were then collected for downstream experiments.

### Immunoprecipitation

Neuro2A cell pellets collected from T-175 flasks (Gen Clone #25-211) at full confluency were lysed in 1 ml NP-40 Lysis Buffer (50 mM Tris-HCl, 150 mM NaCl, 1% NP-40, 1 mM PMSF (Roche #10837091001), 1x cOmplete Protease Inhibitor (Roche #50-100-3301) for 30 minutes on ice and then spun at 16,000 x g for 20 minutes at 4°C and supernatant was transferred to a fresh tube. 25 μL supernatant was then taken as an input control and remaining sample was exposed to either 10 μg of rabbit anti-Arc (Proteintech #16290) antibody or 10 ug of rabbit serum control for 2 hours at room temperature with mixing. Arc protein was then immunoprecipitated using the Pierce protein A/G isolation manual immunoprecipitation protocol (Thermo #88802, Pub No. MANN00157.42). Briefly, 25 μL Pierce Protein A/G Magnetic Beads were cleaned in 1 ml wash buffer and then added to the antibody lysate mixture and incubated at room temperature for 1 hour with mixing. Magnetic beads were then washed in wash buffer and then purified water and Arc protein was then eluted in 100 ul Pierce IgG Elution Buffer (Thermo #21004) with 10 minutes at room temperature with gentle rotation.

### Western Blotting

Extracts from IP experiments, total cell lysate and extracellular vesicle isolations were incubated at 95°C for 5 minutes and resolved by SDS-PAGE in a 4-20% gradient gel (Bio-Rad #4568094) under reducing and denaturing conditions. Proteins were then transferred onto a methanol activated PVDF membrane (Millipore #IPVH00010) and blocked in 5% instant nonfat dry milk in TBST (50 mM Tris (pH 7.4), 150 mM NaCl, 0505% Tween 20) and incubated with primary antibodies diluted in TBST overnight at 4°C. After washing in TBST, blots were incubated with IRDye secondary antibodies (LiCor) for 1 hour at room temperature and imaged using the Chemidoc Touch imaging system (BioRad). The following primary antibodies were used, anti-Mbnl1 (Proteintech # 66837), anti-Arc (Invitrogen #OSA00014G), anti-CD81 (Abclonal #109201), anti-TSG101 (Abclonal 125011), anti-Calnexin (Abclonal #22595), anti-HSP70 (System Biosciences #EXOAB-Hsp70A-1).

### Digital PCR

RNA from IP experiments, total cell lysate and extracellular vesicle isolations was isolated using the RNAeasy mini kit (Qiagen #74104) and then reverse transcribed into cDNA using the AzuraQuant II cDNA Synthesis kit (# AZ-2501) following manufacturer’s protocol using RNAse H digestion. The dPCRs were ran using QIAcuity evergreen master mix (Qiagen 250111) in 8.5K 24-well nanoplates (Qiagen #250011) in a QIAcuity One system. Data was processed in the QIAcuity Software Suite (Qiagen) where absolute values (copies/μl) were obtained and normalized expression derived. The following primers were used for dPCR. Mbnl1: F-CACTGAAAGGTCGTTGCTCCA, R-CGCCCATTTATCTCTAACTGTGT, Arc: F-CACTCTCCCGTGAAGCCATT, R-GGTGCCCACCACATACTGAA, Actin: F-TGGGTGGTCCAGGGTTTCTTACTCCTT, R-ATGTCACGCACGATTTCC.

### Cell Culture Immunofluorescence

Neuro2A were grown on poly-l-lysine coated (Sigma # P8920)glass coverslips (EMS #72222-07) as described in the **Cell Culture** section. Cells were then washed 3 times with PBS and fixed in 4% paraformaldehyde (EMS#15714) for 20 minutes, then washed 3 times in PBS and then permeabilized with 0.2% Triton-X (Sigma #T8787) in PBS for 10 minutes. Cells were then washed 3 times and blocked for 30 minutes in 1% Bovine Serum Albumin, 1% donkey serum in 0.05% Tween-20 in PBS. Cells were then incubated in 1:750 ms anti-tubulin (DSHB #1027) overnight at 4°C. Cells were then washed 3 times in PBS and incubated cells in 1:10,000 Hoechst (Thermo #62249) for 10 minutes and then washed 3 times in PBS. Coverslips were then mounted on slides using Vectashield PLUS Antifade Mounting Media (Vector Laboratories Inc. #H-1900). Images were acquired using a Zeiss LSM 800 (Zeiss, Carl Zeiss NTS Ltd., NY, USA) confocal microscope equipped with a Zeiss 40X Plan-Apochromat 1.30 NA DIC (UV) VIS-IR M27 oil immersion objective.

### Extracellular Vesicle Isolation and Size Distribution Analysis

Media collected off Neuro2A cells was centrifuged at 1000xg for 10 minutes and then filtered using a 0.22 μm filter. Filtered supernatant was then spun at 1500xg for 10 minutes at 4°C and then concentrated to 2 ml in a centricon-70 100 kDa MWCO concentrator (Millipore #UFC710008). Concentrated supernatant was then centrifuged at 10,000xg for 10 minutes at 4°C. Supernatant was then applied to a qEV2 35 nm pore column (Izon #IC2-35) and fractions were collected using the Izon Automated Fraction Collector. Fractions 2-6 were then consolidated and concentrated using a 100 kDa MWCO filter (Millipore #UFC910008) and used for downstream experiments.

Extracellular vesicle size and concentration was then quantified using the Izon Exoid Tuneable Resistive Pulse Sensing system with a NP150 pore with a 46.99 mm stretch, 500 mV voltage and across 800, 1000 and 1400 Pa pressure points and raw data was analysed using the Izon Exoid Data Suite software (version 1.0.2.32).

### Negative Stain Electron Microscopy

EV preparations were fixed overnight in a final concentration of 2% paraformaldehyde (EM grade) at 4°C. After fixation, solution was gently pipetted up and down several times to resuspend EVs. Copper grids coated with carbon film (Electron Microscopy Sciences, CF200-Cu-50) were glow discharged on a PELCO easiGlow at 25 mA for 35 seconds (negative polarity) before use: 10 μL of sample was applied to the grid and incubated for 1 minute. Excess sample was blotted on filter paper, then the grid was rinsed with 15 μL water (filtered by 0.22 μm PVDF membrane) 3 times followed by staining with 1% uranyl acetate (pH 4.5) for 1 minute and then blotted dry. Samples were imaged with a FEI Tecnai Spirit 12 transmission electron microscope at 120 kV equipped with a Gatan 4K camera.

### Mice

All mouse experiments followed the guidelines for care and use of laboratory animals provided by the National Research Council, and with approved animal protocols from the Institutional Animal Care and Use Committee of the University of Massachusetts Chan Medical School (UMCMS). C57Bl/6J (Stock #000664, Jackson), were bred in the UMCMS animal facility and used in mouse experiments as indicated.

### Foot Shock Paradigm

For the foot shock, mice were placed in a 18.5 x 18.5 x 26.6 cm chamber (Ugo Basil) with clear acrylic walls and a metal grid floor through which foot shocks were administered using Anymaze (Stoelting). Foot shock intensity was 0.57 mA for a duration of 2 s, sessions started with a 2 min habituation period during which no shocks were administered. There were 10 foot shock trails with 60 s on average pseudorandom inter-trial-intervals. Mice were left in the in the chamber 30 s after the final shock and then returned to their home cage. Mice were sacrificed one hr after foot shock.

### Dual Fluorescent *in situ* Hybridization and Immunohistochemistry

Sixty min after foot shock, mice were sacrificed, the brains were quickly removed and were snap-frozen in dry ice. A digoxigenin (DIG) 5’ and 3’ -labeled Mbnl1 LNA _TM_ – enhanced oligonucleotide probe was synthesized by Qiagen (Qiagen Sciences, Maryland 20874, USA) designed against the sequence 5’-TGATGCACTGGTGGCTAAGCT-3’ present in the 6^th^ exon of Mbnl1. A LNA _TM_ mRNA detection control probe (ScrambleISH) was purchased from Qiagen as negative control. 20 μm cryostat sections were mounted onto Superfrost Plus slides. Sections were fixed in 4% formaldehyde for 10 min. Following 1 x phosphate-buffered saline (PBS) washes, sections were pretreated with 2 x saline sodium citrate buffer (2 x SSC) for 25 min. Following DEPC water washes, sections were acetylated in 0.1 M triethanolamine (pH 8.0) for 10 dips, then 0.25% (v/v) acetic anhydride for 10 min. Following washes in 2 x SSC, sections were dehydrated in a graded ethanol series (50%, 70%, 95%, 99%) each for 3 min. Hybridizations were performed at 54°C in ISH buffer (sigma) containing probe (200 pmol) overnight. Post-hybridization washes include rinsing in 4 x SSC, 2 x SSC, 0.5 x SSC and 0.1 x SSC, each for 30 min at 58°C. Sections were then incubated in 3% H_2_O_2_ for 15 min and then PBS containing 0.1% Tween 20 for 3 min. Sections were then blocked for 1 h with 1 x PBS/0.3% TritonX-100/5% normal goat serum/1% BSA and then incubated with anti-Arc primary antibody (Proteintech #16290) (1:500) overnight at 4°C in blocking buffer. Following washes with 1 x PBS, sections were incubated with anti DIG-FAB peroxidase (POD) antibody (1:250, Roche) and goat anti-rabbit secondary antibody conjugated to Alexa Fluor 488 (1:800) to reveal the Arc signal for 2 h in a blocking solution containing 0.5% blocking reagent (Roche), 10% heat inactivated goat serum and 0.1% Tween 20. Sections were then washed twice in PBS containing 0.1% Tween 20 followed by applying TSA Plus Cy3 (Perkin Elmer) for visualization of Mbnl1 (FISH). Slides were then washed 3 times in PBS, air dried, and coverslipped with Vectashield mounting medium (Vector Laboratories, Burlingame, CA) were acquired using a Zeiss LSM 800 (Zeiss, Carl Zeiss NTS Ltd., NY, USA) confocal microscope equipped with a Zeiss 40X Plan-Apochromat 1.30 NA DIC (UV) VIS-IR M27 oil immersion objective. Colocalization analysis was performed using the Zen Microscopy Software and quantified using Pearson correlation coefficient (Zeiss, Carl Zeiss NTS Ltd., NY, USA).

### Micrococcal Nuclease Digest

EVs were incubated in either control buffer (2.5 ug/ml Bovine Serum Albumin, 50 mM Tris-HCl pH 8.0, 5 mM CaCl_2_), Micrococcal Nuclease Buffer (50 gel units/mL Micrococcal Nuclease (NEB #M0247S), 2.5 ug/ml Bovine Serum Albumin, 50 mM Tris-HCl pH 8.0, 5 mM CaCl_2_) or Micrococcal Nuclease Buffer supplement with Triton-X (1:20 Triton-X 100 (Sigma #T8787), 50 gel units/mL Micrococcal Nuclease (NEB #M0247S), 2.5 ug/ml Bovine Serum Albumin, 50 mM Tris-HCl pH 8.0, 5 mM CaCl_2_) for 1 hour at 37°C. Nuclease activity was then inhibited using 5 mM EGTA. RNA was then extracted using the RNeasy Mini Kit (Qiagen #74104).

### Fly Husbandry

All flies were raised on low yeast molasses formulation *Drosophila* food at either 25°C or 29°C (Gal4/RNAi crosses).

### Constructs

The GFP-MblA construct was synthesized (Genscript) and using the Gateway system (ThermoFisher), cloned in the pENTR-DTOPO (ThermoFisher) vector and injected into flies and integrated at site attP2 on the third chromosome, through integration by BestGene.

### Larval NMJ Dissections

*Drosophila melanogaster* third instar larva body wall muscles were dissected in calcium-free saline and fixed in either Bouin’s fixative (0.9% (v/v) picric acid, 5% (v/v) glacial acetic acid, 9 % (w/v) formaldehyde) or 4% paraformaldehyde in 0.1 M phosphate buffer, pH 7.2. Fixed samples were washed and permeabilized in PBT (0.1 M phosphate buffer; 0.2% (v/v) Triton X-100 (Sigma #T8787)) and incubated in a primary antibody overnight at 4°C. The samples were then washed 3 times with 1x phosphate-buffer saline (PBS) with 0.05% Tween-20 (PBT), incubated with secondary antibodies for 2 hours at room temperature, washed 3 times and mounted in Vectashield PLUS Antifade Mounting Media (Vector Laboratories Inc. #H-1900). The following antibodies were used: rabbit anti-DLG, 1:40,000 [28], goat anti-HRP DyLight 405 (Jackson ImmunoResearch #123-475-021), 1:200, rabbit anti-MblA, 1:500 [13], mouse anti-GFP (DSHB # 4C9), 1:500, rabbit anti-dArc1 [7], 1:500, Hoechst (Thermo #33342), 1:50,000. DyLight-conjugated and Alexa Fluor-conjugated secondary antibodies were obtained from Jackson ImmunoResearch (Alexa Fluor-594-congugated goat anti-HRP, Alexa Fluor-488-congugated donkey anti-Rabbit, Alexa Fluor-594-congugated goat anti-rabbit, Alexa Fluor-647-congugated goat anti-Mouse) and were used at 1:200, as described above.

### Confocal Microscopy and Signal Intensity Measurements

Z-stacked images were acquired using a Zeiss LSM 800 confocal microscope equipped with a Zeiss 63X Plan-Apochromat 1.40 NA DIC M27 oil immersion objective and a Zeiss 40X Plan-Apochromat 1.30 NA DIC (UV) VIS-IR M27 oil immersion objective. After image acquisition with identical settings, the images were quantified as previously described [29]. In brief, volumetric measurements of the boutons of interest bound by HRP staining were selected and fluorescence intensity inside was measured using Volocity software (Quorum Technologies Inc.). Briefly, after image acquisition, the bouton volume bounded by HRP staining was selected using Volocity, and fluorescence intensity inside was measured. To calculate the postsynaptic area, the presynaptic bouton selection was dilated by 6 iterations, the HRP containing volume was subtracted and the intensity within the remaining volume was measured. Intensity was determined as the sum of total pixel intensity in each volume (pre- and postsynaptic) and normalized to bouton volume, as described previously [29].

### Quantification and Statistical Analysis

Statistical analyses for single comparisons were performed using a Student’s *t* test while multiple comparisons with experimental groups utilized a one-way analysis of variance (ANOVA) with the appropriate post hoc test. *, *p* < 0.05; **, *p* < 0.001; ***, *p* < 0.0001. Raw data files were processed with Excel (Microsoft) and data analysis for statistical significance utilized GraphPad Prism version 9.5.0 (GraphPad Software).

## Acknowledgements

We appreciate the Bloomington Drosophila Stock Center, FlyBase and the Vienna Drosophila Resource (VRDC) for invaluable reagents and resources. We thank Alfred Simkin and Gimena Alegre for valuable discussions on computational analysis. This work was supported by National Institute on Drug Abuse, DA041482 (A.R.T.); and DA047678 (A.R.T.) and National Institute of Neurological Disorders and Stroke, RO1NS112492 to T.T.

## Figures

**Supplementary 1.**
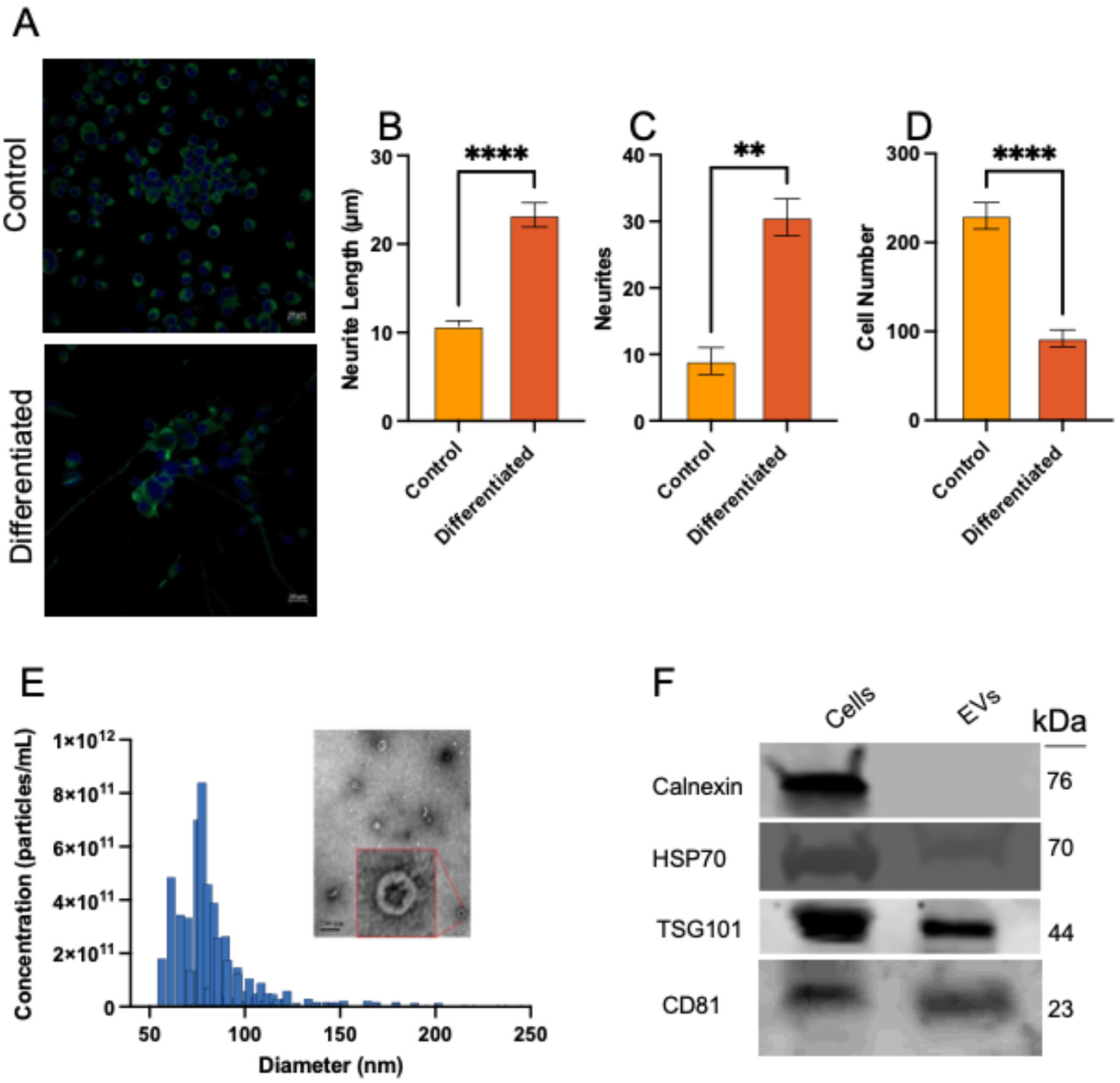
**A**. Micrographs of control and differentiated Neuro2A cultures (20 μm scale bar) showing **B**. increased neurite length **C**. increased neurite number and **D**. reduced cell division. Error bars, mean ± SEM; Statistical analysis conducted using an unpaired T-test. **E**. Size and concentration analysis of N2A EVs by TRPS. Inset: Negative stain electron micrograph of N2A EVs **F**. Western blot analysis of Neuro2A derived EVs and cell lysate. Error bars, mean ± SEM; Statistical analysis conducted using an unpaired T test; P values: *<0.05, NS, not significant.

**Supplementary 2.**
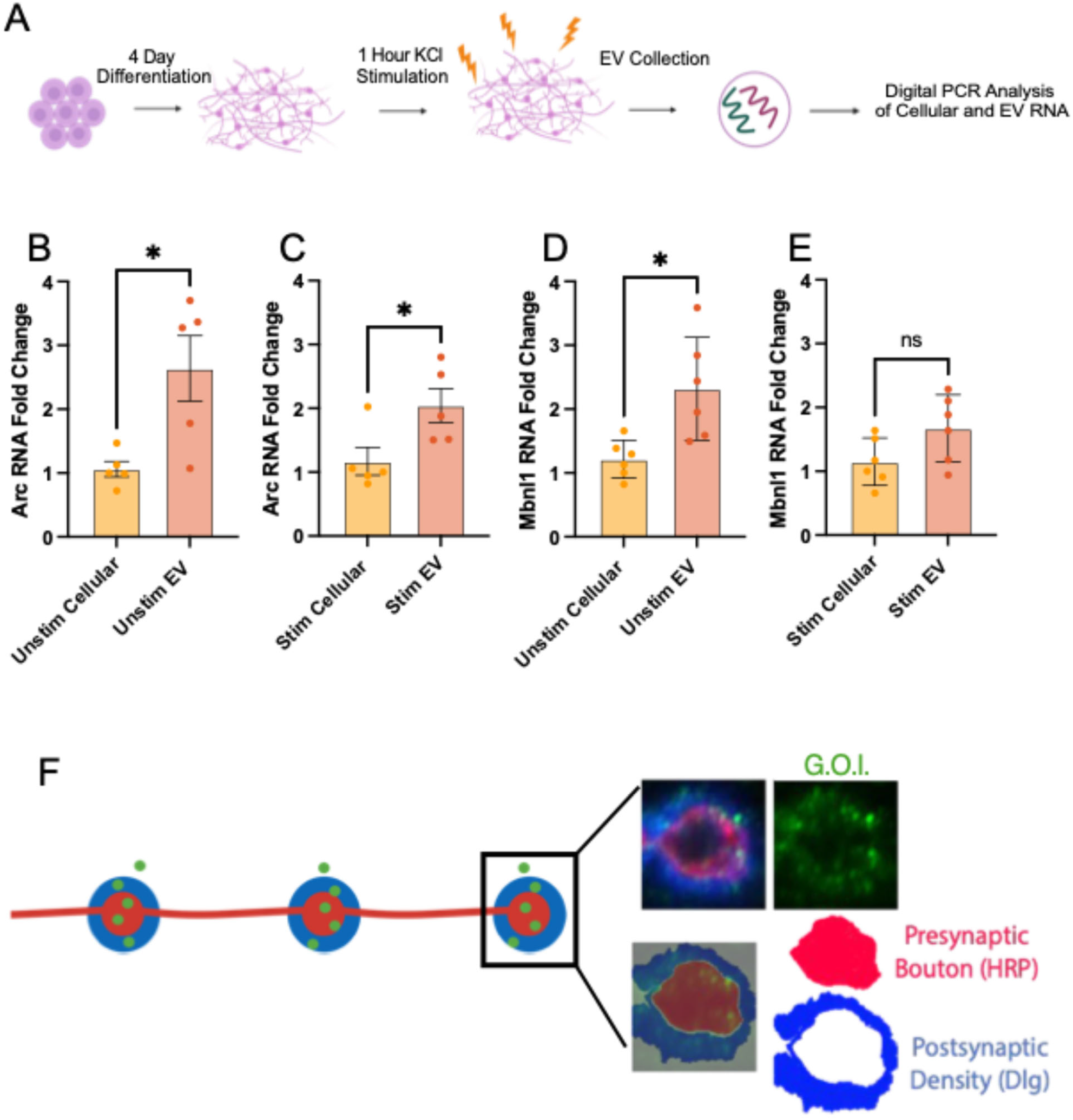
**A**. Experimental workflow schematic. Neuro2A were differentiated in reduced serum media and retinoic acid for four days and then stimulated with KCl for 1 hour prior to EV isolation. **B**. dPCR detecting unstimulated differentiated Neuro2A EV Arc compared to cellular Arc. **C**. dPCR detecting stimulated differentiated Neuro2A EV Arc compared to cellular Arc. **D**. dPCR detecting unstimulated differentiated Neuro2A EV Mbnl1 compared to cellular Mbnl1. **E**. dPCR detecting stimulated differentiated Neuro2A EV Mbnl1 compared to cellular Mbnl1. **F**. Schematic demonstrating the anatomy and compartmentalization of the *Drosophila* larval NMJ. Error bars, mean ± SEM; Statistical analysis conducted using an unpaired T test; P values: *<0.05, NS, not significant.

